# The neural signature of simple overlearned temporal expectations resembles episodic retrieval

**DOI:** 10.64898/2026.07.07.737020

**Authors:** Linda Sempf, Peter Vavra, Toemme Noesselt

## Abstract

Temporal expectations are crucial for adaptive responding to predictable events. Most previous imaging studies have focused on the neural underpinnings of unexpectedness rather than expectedness, without considering that neural responses related to temporal expectations may be confounded by parallel learning processes. To identify the neural basis of overlearned expectancy with fMRI, we employed a multi-day detection paradigm with auditory cues preceding visual targets at likely and unlikely time points. After training left inferior parietal cortex was engaged in processing overlearned temporal expectations together with retrosplenial and prefrontal cortex. Remarkably, compared with untrained controls, we observed inverted brain responses as a function of stimulus likelihood and training status. Finally, the brain regions coding overlearned expectancy had previously been implicated in episodic retrieval and simulation (based on Neurosynth association tests), suggesting that frequently used temporal expectations may, over time, utilize episodic memory routines to simulate likely future events and thus guide behavior.

## Introduction

Our environment is rich in statistical regularities that convey meaningful information especially within stable environments. For instance, we learn that the brief whir of an automatic door motor signals that the doors to the cafeteria will open a certain moment later, and we routinely use this information to adjust our walking speed without risking a collision or wasting time waiting in front of a still-closed door. The cognitive system is well adapted to detect temporal regularities, as this capacity allows us to foresee possible consequences and prepare those that appear most likely^1^. This skill might be particularly useful in reoccurring environments, such as those encountered in the course of our often stubborn daily work routines. A large body of empirical research suggests that we continuously rely on temporal expectations to optimize motor responses (for a comprehensive review, see^2^). For instance, in simple detection paradigms temporal predictability reliably facilitates response speed, resulting in faster reactions to expected stimuli: the so-called temporal expectation effect.

These temporal expectations are thought to be the product of learning of temporal regularities of the environment. The cognitive system continuously extracts temporal structures from incoming sensory information, which can then be used to form new expectations. In probabilistic environments this extraction process is commonly described by statistical learning, i.e. an innate, automatic function of the cognitive system^3,4^ and has been related to specific processes including updating and surprise. Several hypotheses have been proposed regarding the neural basis of this learning phenomenon. In a recent review, ^5^ noted that statistical learning ‘includes practically the entire brain’, reflecting the fact that domain-general processes, domain-specific processes, and their interaction have all been proposed as explanatory accounts. Such domain-general computational principles may be supported by partially shared neural systems, including the hippocampus, whereas domain-specific representations formed through statistical learning remain constrained by the sensory and motor systems processing the relevant information^6^.

Another line of research focuses on the effects of utilizing temporal expectations which are often induced through statistical regularities embedded in the experimental design. Within these implicit temporal expectation paradigms, involved brain regions include prefrontal regions plus left fronto-cingulate cortex^9^, motor regions including SMA, premotor cortex, and cerebellum^14^, temporal regions including posterior middle temporal gyrus^15^, superior temporal gyrus^12^, planum temporale^16^ and even primary auditory cortex^7^, and occipital cortex including even primary visual cortex^16^. Most prominently, the left inferior parietal cortex, and left intraparietal sulcus have been repeatedly associated with the active application of temporal expectations and even more fundamental time–space integration processes^20^ - though these activations often emerge in task-related contrasts (e.g., temporal attention versus no temporal attention or exogenous versus endogenous timing tasks), and the exact location within parietal cortex and sometimes even the lateralization vary between studies. Taken together, previous research points to a wide range of neural regions implicated in the application of temporal expectations, partially overlapping with regions associated with statistical learning. Yet, the differentiation between processes related to the dynamical formation versus the active use of already formed expectations has so far been neglected, as most studies investigating the application of temporal expectations typically assess these expectations in the same session in which the underlying regularities are acquired, and therefore after only limited exposure to these temporal regularities, such that their formation and use are likely to occur concurrently.

On a theoretical level, Bayesian accounts of learning^21–23^ and related predictive coding frameworks^24–27^ all posit that learning is strongest during the initial stages of exposure and decreases over time. Likewise, as the environmental structure becomes familiar and predictions become more reliable, learning gradually saturates and updates to the internal model become smaller. First evidence for this proposed interaction of acquisition and application in temporal expectation was reported by ^8^, though note that this study examined the dynamics only within a single session, whereas in real-world settings temporal expectations are often repeatedly retrieved and applied over much longer time scales and may be further consolidated during sleep. Hence, repeated exposure over days should minimize the effects of learning on brain responses.

### The current experiment

Here, we aimed at identifying neural regions instrumental in the application of temporal expectation in stable environments. By the repeated exposure prior to our main experiment, we aimed to minimize learning and updating. To mirror real-life as closely as possible, passive observations and active task engagement were both introduced in the experimental design: an initial passive viewing phase resembling statistical learning through observation^28,29^, followed by a two-day training program requiring participants to actively use the temporal regularities to optimize their performance. Only after these three exposure phases, on the fourth day of the study, did we examine the neural underpinnings of the application of overlearned temporal expectation (see Fig. 1a).

**Fig. 1.**
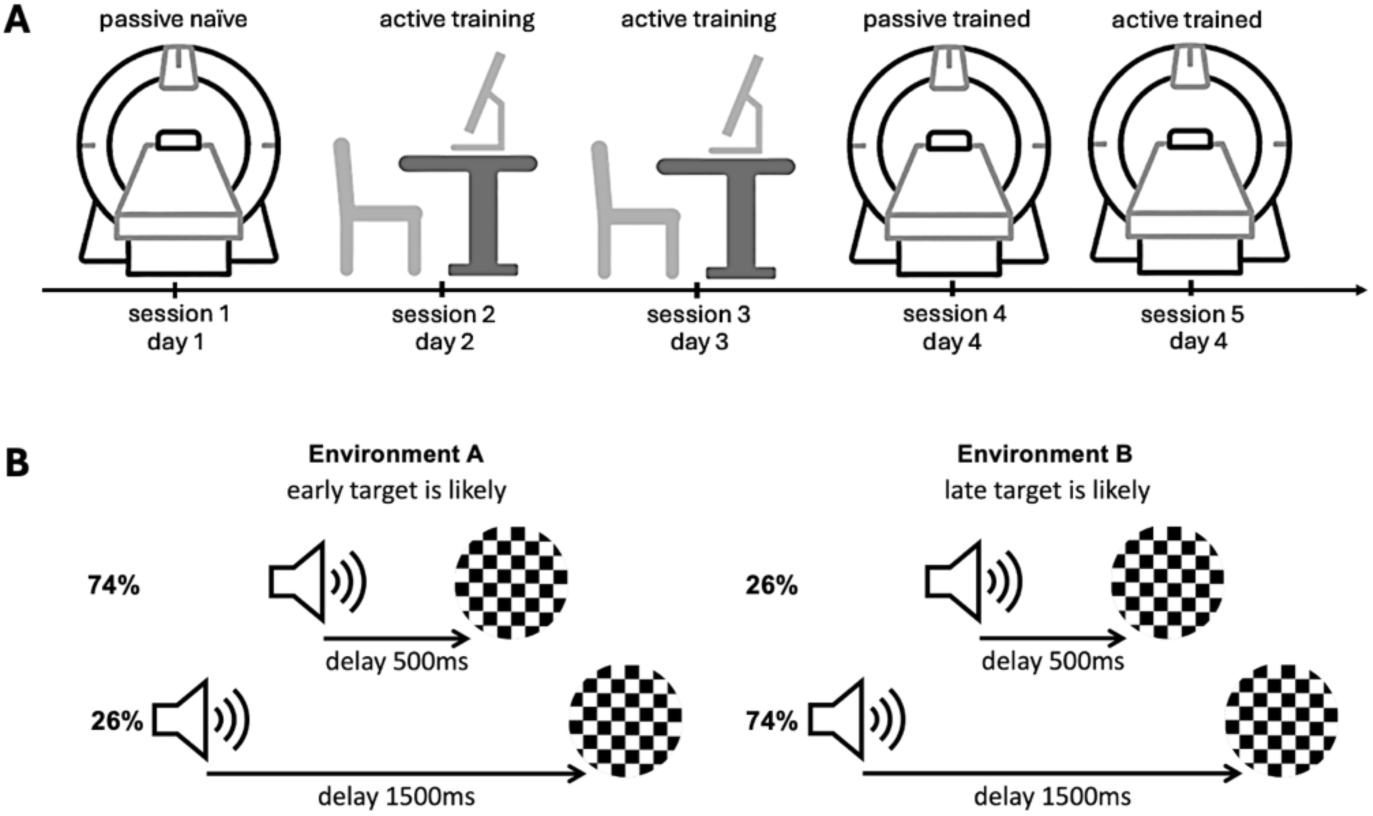
**A** Experimental procedure. In the main experiment participants completed five sessions across four consecutive days: a passive naïve session in the scanner (session 1, day 1), two active training sessions outside the scanner (session 2–3, days 2-3), a passive trained session in the scanner (session 4, day 4), and an active trained session in the scanner (session 5, day 4. In a control experiment, participants only completed one active session inside or outside the scanner (identical to session 5 or 2 but without prior training or passive exposure). **B** Experimental design with two different temporal environments. In environment A (early target likely), the visual target occurred 500 ms after the auditory cue on 74% of trials and after 1500 ms on 26%. In environment B (late target likely), the probabilities were reversed, with the target appearing after 1500 ms on 74% and after 500 ms on 26% of trials.

In particular, we designed a study in which participants were confronted with cross-modal stimulus sequences, enabling the formation of distinct temporal expectations (see Fig. 1b). The task design consisted of an auditory cue followed by a visual target appearing either after 500 or 1500 ms. Within blocks of stimulus sequences, targets were either likely or unlikely to occur at an early or late time point (74%/26% likelihood ratio of the two cue-target intervals). Thus, there were two distinct temporal environments: one in which the target appeared early on most trials (*early likely block*), and one in which the target appeared late on most trials (*late likely block*) – see Fig. 1b.

Behavioral and fMRI read-outs were recorded and we examined how temporal expectation effects – that is, improved performance in response to temporally likely vs unlikely stimuli – are reflected in overt motor behavior and brain signals. These overlearned read-outs were compared to those acquired without training. In particular, we hypothesized that *early likely* trials should rely on the application of temporal expectation, whereas *early unlikely* trials should rely on a violation of temporal expectation, often termed ‘surprise’. The *late likely* trials should again rely on cognitive process of expectation application. However, *late unlikely* trials would not only rely on surprise but a readjustment of the temporal focus of attention to another point in time in contrast to unlikely early trials. Hence, the comparison of *early likely* vs *unlikely* trials should reveal the effects of temporal expectation less contaminated by additional cognitive processes.

## Results

### Temporal regularities shape response times

We first compared *early likely* to *early unlikely* trials behaviorally. On average, participants responded significantly faster to *early likely* compared to *early unlikely* trials (t(34) = -6.66; p < .001; see Fig. 2a and table 1). This suggests that participants were able to build temporal expectations based on temporal regularities - a notion consistent with the idea that anticipating events in time facilitates responses (e.g.,^30^).

**Fig. 2.**
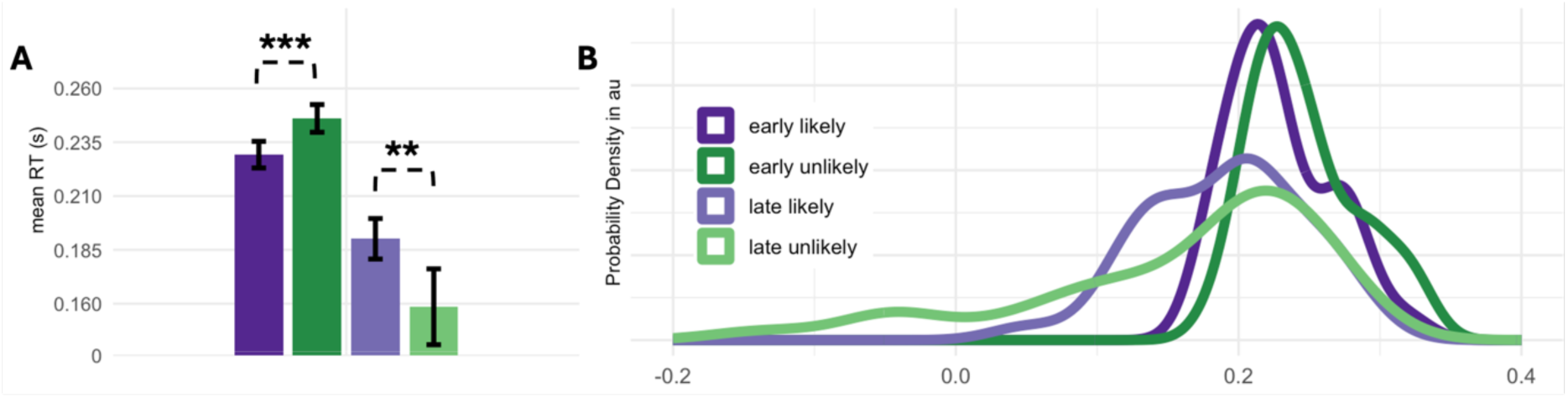
Group average response times (RT) for *early likely* targets are displayed in dark purple and for *early unlikely* targets in dark green, whereas response times for *late likely* targets are shown in light purple and for *late unlikely* targets in light green. **(A)** Mean RTs (± SE). Participants responded faster to *early likely* than to *early unlikely* targets. In contrast, participants responded faster *to late unlikely* than to *late likely* targets, reflecting premature responses resulting in very short and, in some cases, even negative RTs. Note that this inverse pattern remains even if premature responses (i.e. < 200 ms post-target) were excluded from analysis (p<.057). Also note that the use of median RTs for statistics resulted in qualitatively similar outcomes. Statistical significance is denoted by *** for early trials and ** for late trials. **(B)** Density distribution of mean RTs across participants for all four conditions. Density distributions illustrate lower RTs *for early likely* compared to *early unlikely* targets and the presence of very short and, in some cases, even negative RTs in late trials. 0 denotes target onset. Y-axis represents estimated probability density in arbitrary units.

**Table 1.** Mean (M) response times (ms) and standard error (SE) for each condition, collapsed across all runs and separately for each run.

|  | Early likely | Early unlikely | Late likely | Late unlikely |
| --- | --- | --- | --- | --- |
| Overall | 229.31 (6.22) | 245.66 (6.39) | 189.93 (9.45) | 158.90 (17.53) |
| Run 1 | 230.04 (6.82) | 244.33 (9.37) | 199.73 (9.40) | 180.02 (14.94) |
| Run 2 | 231.82 (6.39) | 239.93 (5.87) | 195.60 (10.02) | 157.79 (22.00) |
| Run 3 | 225.30 (6.41) | 249.73 (8.59) | 196.61 (11.66) | 143.13 (20.50) |
| Run 4 | 230.08 (7.74) | 248.66 (7.83) | 167.78 (14.36) | 154.65 (22.85) |

Note. Values are reported as M (SE).

No consistent correlate of expectancy has been proposed for *late likely* vs. *unlikely* trials (though see^31,32^), and it has further been argued that expectancy is diminished for late trials^33^. Here, we observed even negative response times in numerous late trials (see Fig. 2b), resulting in a “negative” temporal expectation effect, with shorter response times to *late unlikely* than to *late likely* targets. This may indicate that participants also formed temporal expectations for late trials. However, the pattern observed for late trials may partly reflect anticipatory responses triggered by expectations at the early time point. In *late unlikely* trials premature responses may reflect a strong anticipation of target occurrence at the early time point due to the predominance of early targets within that environment. Enhanced response preparation, combined with failed response inhibition, may therefore have resulted in a higher number of premature responses here. In *late likely* trials premature responses may reflect increased uncertainty due to the longest cue-target interval (see e.g.^34,35^) resulting in an increase in premature responses with increasing cue-target intervals^31^.

Next, we examined the statistical robustness and time course of these temporal expectation effects for early trials. The effect of temporal expectation remained significant (main effect likeliness: F(1,34) = 38.31, p <.001) and did not vary across runs (interaction run x likeliness: F(2.19, 74.44) = 1.41, p = .249), see Fig. 2a. For late trials we again observed a main effect of likeliness (F(1,34) = 9.43, p <.005) and no significant interaction with run F(2.30, 78.25) = 1.64, p = .197). In sum, participants showed a robust temporal expectation effect, though inverted for late trials.

In the two combined control experiments, we again found that naïve (i.e., untrained) participants responded significantly faster to *early likely* than to *early unlikely* targets (main effect likeliness: F(1,50) = 139.90, p < .001) and temporal expectation effects did not vary across runs (interaction run x likeliness: F(2.67, 133.71) = 0.09, p = .954). Also consistent with the main experiment, late trials showed a main effect of likeliness (F(1, 50) = 17.39, p < .001), with faster response times for *late unlikely* than for *late likely* trials. This effect did not interact with run (F(1.82, 90.77) = 0.87, p = .413).

Finally, we directly tested whether the repeated exposure to the stimuli modulated behavioral read-outs of temporal expectation relative to a single-session version of the task. To this end, response times for early trials of trained and naïve participants were compared using a linear mixed-effects model. As previously observed, temporal expectation effects were significant (main effect likeliness: F(1, 17010.07) = 162.78, p < .001). There was neither a main effect of training status (F(1, 83.66) = 0.00, p = .993), nor a main effect of experimental setting (inside/outside MRI-scanner; F(1, 82.98) = 0.42, p = .519), indicating comparable overall response times that were unaffected by general training effects or by the MRI environment. Importantly, the interaction between likeliness and training status was significant (F(1, 17010.07) = 11.17, p = .001), with a significantly larger TE effect in naïve than in trained participants (Δ = 12.6 ms, z = -3.34, p < .001). No such differential interaction effect was observed for late trials (p = .534). This indicated that repeated exposure and training indeed altered the behavioral expression of temporal expectations selectively for early trials.

### Neural correlates of overlearned temporal expectation resemble those of episodic cognition

Following this, we aimed to identify the brain regions instrumental in the application of overlearned temporal expectation. First, we characterized the neural basis of the temporal expectation effect for early trials resembling the behavioral effect and thus facilitating comparability with previous studies investigating temporal expectations (e.g. ^36^). Comparing *early likely* > *early unlikely* trials, two significant regions emerged: one in the left inferior parietal lobule (IPL) and one in the left rostrolateral prefrontal cortex (RLPFC) (see Fig. 3a, yellow areas and table 2). Moreover, statistical regularities might still have been extracted in both early and late temporal environments, while surprise and attentional reallocation might have overshadowed behavioral readouts of temporal expectation in the late conditions. To test this hypothesis of a general expectancy effect on the neural level, we therefore compared *likely > unlikely* trials across temporal environments. We observed significant modulations of fMRI signals spanning the left IPL, the anterior retrosplenial cortex (RSC), right anterior medial prefrontal cortex (aMPFC), and bilateral ventromedial prefrontal cortex (vmPFC) (see Fig. 3a, green areas and table 2). For completeness, we also tested for additional regions significantly modulated by *late likely > unlikely trials*; however, no clusters survived correction for multiple comparisons. Bar plots of the region-wise average beta weights revealed a difference for both early and late trials (though note that the difference of late trials were slightly smaller than of the early trials, see Fig. 3b). Together, these results indicate that a network of brain regions is instrumental in successfully applying overlearned temporal expectations and may account, at least in part, for the observed behavioral temporal expectation effect.

**Fig. 3.**
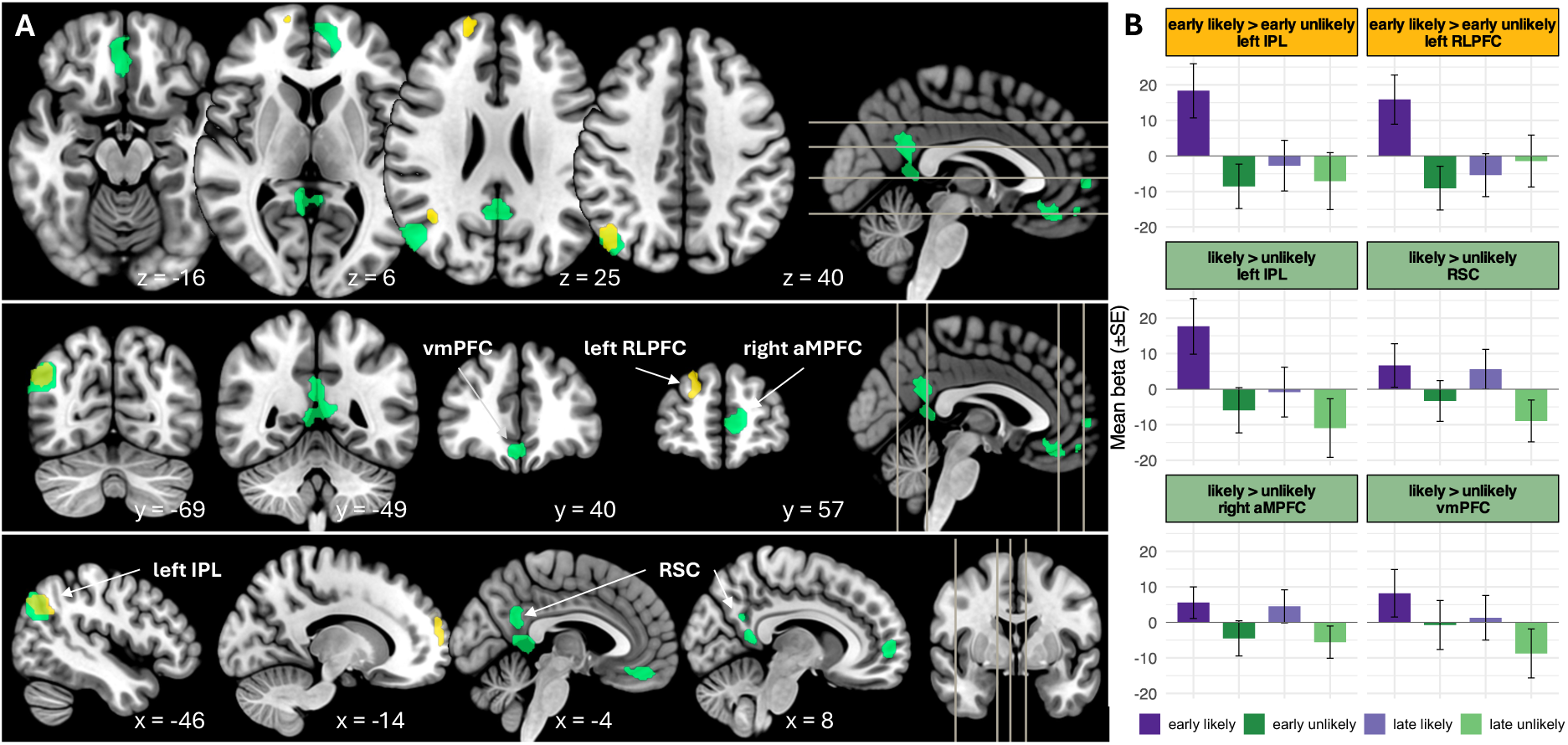
**(A)** Neural correlates of overlearned temporal expectation. Whole-brain group results for the main effect of temporal likeliness (*likely > unlikely*) are displayed in green, the effect of temporal expectation in early trials (*early likely > early unlikely*) is displayed in yellow. For early trials, significant clusters were observed in the left inferior parietal lobule (left IPL, BA 39) and the left rostrolateral prefrontal cortex (left RLPFC BA 10). For the main effect (likely > unlikely), significant clusters were observed in the left inferior parietal lobule (left IPL, BA 39), retrosplenial cortex (RSC, BA 29/30), right anterior medial prefrontal cortex (right aMPFC, BA 10), and ventromedial prefrontal cortex (vmPFC, BA 11). Statistical maps are displayed on a standard Montreal Neurological Institute template (MNI152) and thresholded at p < .05 (FWE-corrected at the cluster level; auxiliary threshold of p < .001 at the voxel level). Color-coded regions indicate enhanced fMRI-signals for likely compared to unlikely trials. Coordinates are reported in MNI space. X-, Y-, and Z-values indicate slice location in MNI space. Neurological convention is used. **(B)** Region-wise mean beta weights (proportional to % signal change) for each cluster.

**Table 2.**
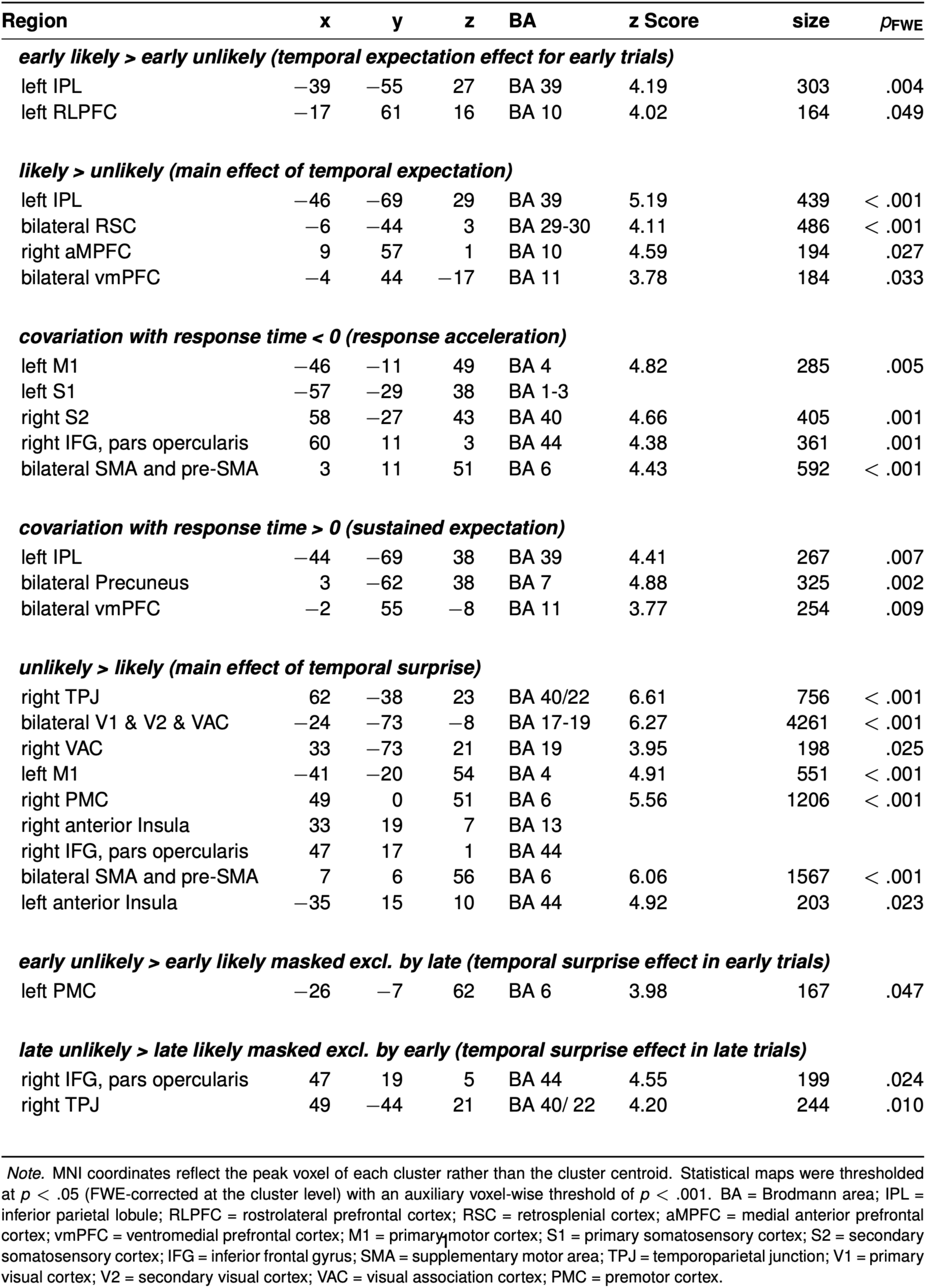
Whole-brain group results for temporal expectation, temporal surprise, and response time effects.

### Linking neural activation patterns to cognitive terminology

To further characterize the functional profile of the regions identified in the *likely >* unlikely contrast, we queried the peak voxel of each significant cluster using the term-based association maps provided by Neurosynth^37^. The peak voxels of the left IPL and the RSC showed strong associations with the term episodic (left IPL: z = 7.56, posterior probability = .75; RSC: z = 4.68, posterior probability = .75) and autobiographical (left IPL: z = 9.2, posterior probability = .84; RSC: z = 5.25, posterior probability = .83). The peak voxel of the left IPL showed furthermore strong associations with the term retrieval (z = 7.96, posterior probability = .72). The peak voxel of the aMPFC was strongly associated with the terms default (z = 6.81, posterior probability = .74), autobiographical memory (z = 5.18, posterior probability = .84) and referential (z = 4.3, posterior probability = .77), whereas the peak voxel of the vmPFC was strongly associated with the terms mentalizing (z = 7.65, posterior probability = .84) and theory of mind (z = 5.64, posterior probability = .80).

To complement this voxel-wise characterization, we additionally decoded the entire unthresholded second-level statistical map using the Neurosynth Decoder. Among the highest-ranking terms associated with the resulting activation pattern were default mode (r = 0.267), autobiographical (r = .29), episodic (r = .225), retrieval (r = .195), memories (r = .18), referential (r = .167), self-referential (r = .166), theory of mind (r = .159), recollection (r = .151), mentalizing (r = .146), remembering (r = .144), and construction (r = .125); terms that conceptually converge on the broader construct of episodic cognition^38,39^ or “Mental Time Travel”^40^.

### Response acceleration

To directly test for the correspondance of behavioral and neural measures, and to dissociate temporal expectation signals from simple response time variations, we also identified brain regions in which neural signals covaried with subjects’ response times across conditions. fMRI signals in four clusters showed a negative covariation with response times: first, a cluster spreading from left primary motor cortex to left primary somatosensory cortex, a second cluster within right secondary somatosensory cortex, a third cluster within the pars opercularis region of the right inferior frontal gyrus, and finally, a cluster encompassing the SMA and pre-SMA (see Fig. 4, blue areas and Table 2). This indicates that response acceleration was mainly associated with activity in motor-related regions and not the regions related to overlearned temporal expectation reported above.

**Fig. 4.**
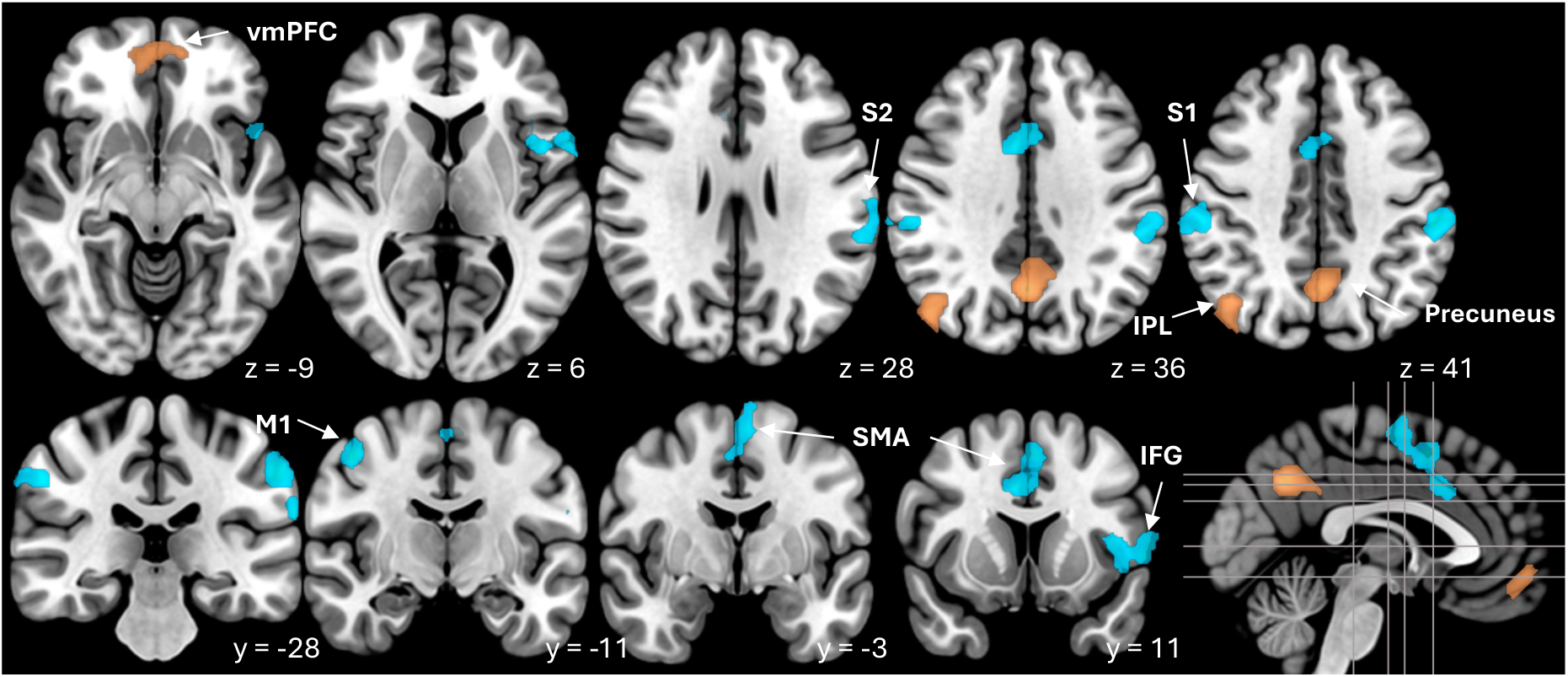
Neural covariations with response time. Whole-brain group results for negative covariation of fMRI-signals with response times (response acceleration) are displayed in blue, results for positive covariation of fMRI-signals with response time (sustained expectation) are displayed in copper. For response acceleration significant clusters were observed in left primary motor cortex (left M1, BA 4), left primary sensory cortex (left S1, BA 1/2/3), right secondary sensory cortex (right S2, BA 40), bilateral SMA and pre-SMA (BA 6) and right inferior frontal gyrus, pars opercularis (right IFG, BA 44). For sustained expectation significant clusters were observed in left inferior parietal lobule (left IPL, BA 39), bilateral precuneus (BA 7) and ventromedial prefrontal cortex (vmPFC, BA 11). Statistical maps are displayed on a standard MNI template (MNI152) and thresholded at p < .05 (FWE-corrected at the cluster level; auxiliary threshold of p < .001 at the voxel level). Color-coded regions indicate enhanced fMRI-signals covarying with slower or faster response times. Coordinates are reported in MNI space. Y- and Z-values indicate slice location in MNI space. Neurological convention is used.

### Sustained expectation

In addition, we identified regions whose fMRI-signals increased with longer response times using the inverse comparison (positive correlations of fMRI-signals with response time across conditions). Here, three significant clusters were found, including the bilateral precuneus, bilateral vmPFC, and once more the left IPL (see Fig. 4, copper areas and Table 2). Figure 5 depicts the three partially overlapping left IPL clusters identified across the comparisons *early likely > early unlikely*, *likely > unlikely*, and the covariation of fMRI-signals with response times. In the context of a robust temporal expectation effect across participants, positive correlations with individual response time should, however, be interpreted carefully. Due to the occurrence of very short and even negative response times, larger mean response times do not necessarily reflect weaker temporal expectations, but may rather reflect a reduced tendency to respond prematurely before target onset. Accordingly, fMRI-signals positively associated with response times may index neural processes related to the maintenance of temporal expectation until the actual target presentation, rather than the formation of temporal expectations per se. In accord, some researchers described the IPL as an episodic buffer^41^.

**Fig. 5.**
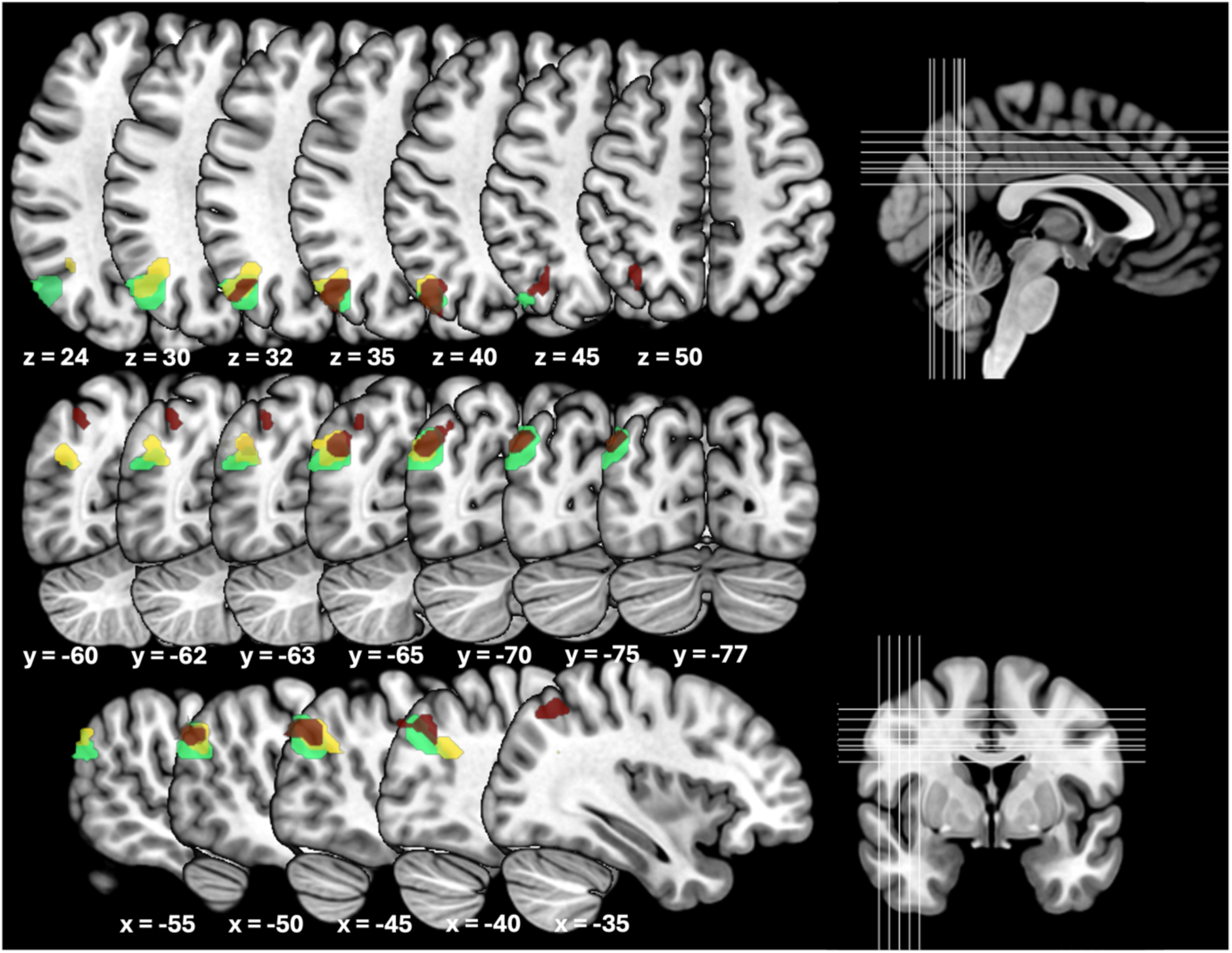
Spatial overlap and dissociation of activation patterns within the left inferior parietal lobule (IPL) across comparisons. Green indicates the main effect of temporal expectancy (*likely > unlikely*), yellow areas indicates the effect of temporal expectancy selectively in early trials (*early likely > early unlikely*), and brown indicates positive covariations of fMRI-signals with slower condition-specific response times. The visualization highlights that all three effects were localized within the left IPL while forming spatially distinct and only partially overlapping clusters. Statistical maps are displayed on a standard MNI template (MNI152) and thresholded at p < .05 (FWE-corrected at the cluster level; auxiliary threshold of p < .001 at the voxel level). Coordinates are reported in MNI space. X-, Y-, and Z-values indicate slice location in MNI space. Neurological convention is used.

### Effect of Temporal Unexpectedness

As a manipulation check we reversed the direction of the main effect of likeliness and examined regions showing enhanced fMRI signals for unlikely relative to likely targets. As expected, the GLM revealed significant effects in seven clusters potentially related to temporal reallocation and surprise: (1) the right temporoparietal junction, (2) bilateral primary, secondary and visual association cortex, (3) an additional cluster within the right visual association cortex, (4) the left primary motor cortex, (5) a cluster encompassing the right premotor cortex, the right anterior insula, the pars opercularis regions of the right inferior frontal gyrus, (6) bilateral SMA and pre-SMA, and (7) the left anterior insula (see Fig. 6 red areas and table 2). Remarkably, the identified clusters closely match the combination of multimodal and unimodal visual networks for involuntary attention to changes in the sensory environment described by ^42^ and, more recently, in attentional reorienting following violations of cross-modal expectancies ^43^. Restricting the analysis to early trials, the comparison *early unlikely > early likely* masked excl. by *late unlikely* > *late likely* at p = .05 revealed a single significant cluster in the left premotor cortex. In contrast, the corresponding comparison for late trials (*late unlikely > late likely* masked excl. by *early unlikely* > *early likely* at p = .05) yielded significant clusters in the right inferior frontal gyrus and the right TPJ (also see Fig. 6 yellow (early) and blue (late) areas, and table 2) pointing at an involvement of these regions in attentional reallocation and surprise as hypothesized above.

**Fig. 6.**
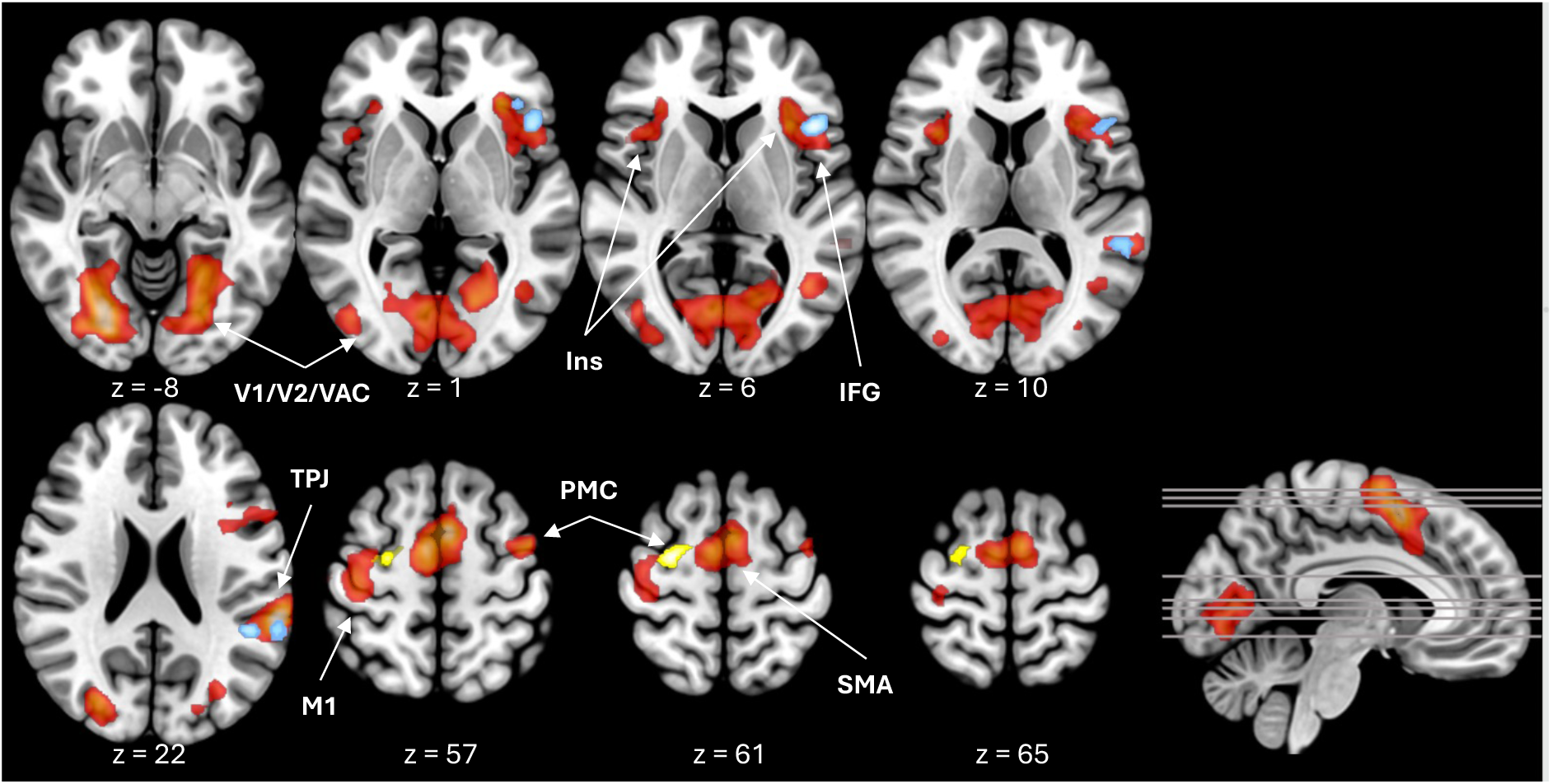
Neural basis of temporal unexpectedness. Whole-brain group results for the main effect of *unlikely > likely* are shown in red.). Results specific to early trials (*early unlikely* > *early likely*, exclusively masked by *late unlikely* > *late likely* at p = .05) are shown in yellow, whereas results specific to late trials (*late unlikely* > *late likely*, exclusively masked by *early unlikely* > *early likely* at p = .05) are shown in blue. For the main effect, significant clusters were observed in the right temporoparietal junction (TPJ), visual cortex (V1/V2) and visual association cortex (VAC), primary motor cortex (M1), supplementary motor cortex (SMA/pre-SMA), and right inferior frontal cortex (IFG, see also Table 2). The early-trial surprise effect was confined to the left premotor cortex, whereas the late-trial effect involved the right inferior frontal gyrus and right TPJ. Statistical maps are displayed on a standard MNI template (MNI152) and thresholded at p < .05 (FWE-corrected at the cluster level; auxiliary threshold of p < .001 at the voxel level). Color-coded regions indicate enhanced fMRI-response for unlikely trials. Coordinates are reported in MNI space. X-, Y-, and Z-values indicate slice location in MNI space. Neurological convention is used.

### Naive control group

Finally, we directly tested for differential effects of training on the neural signature of the temporal expectation effect. We extracted condition-wise beta weights in the trained group (main experiment) and statistically compared the effects with the naive group per region.

We found a significant interaction of likeliness and training status in RSC (F(1,49) = 11.20, p = .002) and right aMPFC (F(1,49) = 6.83, p = .012). Post hoc comparisons showed significant expectancy modulations in trained participants (RSC: p < .001; right aMPFC: p < .001), but not in untrained participants (RSC: p = .634; aMPFC: p = .785; Fig. 7A). No corresponding significant interaction was observed in left RLPFC, neither for the interaction of likeliness and training status (p = .08) nor for the interaction of likeliness, time interval and training status (p = .088), though a trend was there.

**Fig. 7.**
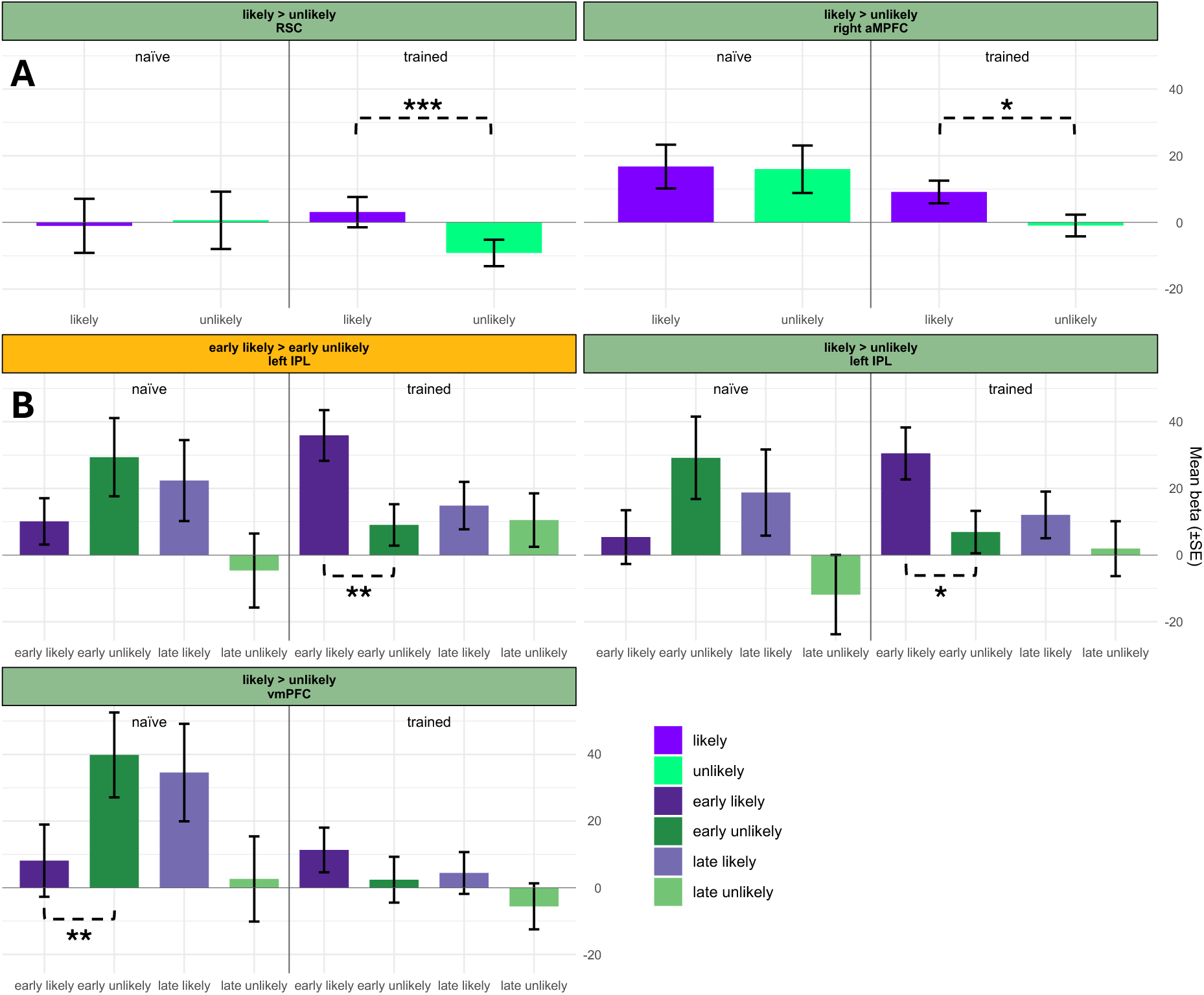
Effects of training on beta weights (proportional to % signal change) within brain region identified in the main experiment. **(A)** Regions showing a significant likeliness × training status interaction. In both RSC and right aMPFC, expectancy-related activity differed between naïve and trained participants. Green panels depict effects based on the overall main effect of likeliness (likely > unlikely), collapsed across cue-target intervals. **(B)** Regions showing a significant likeliness × time interval × training status interaction. Follow-up analyses indicated that these effects were primarily driven by the early cue-target interval, with training modulating expectancy-related responses. Yellow panels depict contrasts restricted to *early likely* > *early unlikely* trials. Bars represent mean beta estimates ± SE.

In left IPL and vmPFC, a significant three-way interaction between likeliness, time interval, and training status was observed (left IPL 1 (region with maximal response to early effect): F(1, 49) = 4.53, p = .038; left IPL 2 (region with maximal response to main effect of likeliness): F(1, 49) = 4.36, p = .042; vmPFC: F(1, 49) = 4.98, p = .030). Follow-up analyses restricted to early trials confirmed that the effect of likeliness differed as a function of training status (left IPL 1: F(1, 49) = 7.49, p = .009; left IPL 2: F(1, 49) = 7.60, p = .008; vmPFC: F(1, 49) = 8.49, p = .005). Pairwise comparisons revealed a significant temporal expectation effect for early trials in trained participants in left IPL (left IPL 1: t(49) = 2.846, p = .006; left IPL 2: t(49) = 2.452, p = .018), whereas no expectancy-modulation was observed in untrained participants (left IPL 1: t(49) = -1.379, p = .174; left IPL 2: t(49) = -1.671, p = .101). In vmPFC, no significant expectancy effect was observed in trained participants for early trials (t(49) = 1.14, p = .259), whereas untrained participants showed a significant reversed expectancy effect favoring *early unlikely* over *early likely* targets (t(49) = -2.75, p = .008). This pattern of results strongly suggests that training has a profound effect on neural representations of temporal expectancy.

## Discussion

In this study, we aimed at identifying the neural underpinnings of the use of overlearned temporal expectations in humans. To this end, we created two distinct temporal environments which differed only in the probability distributions of cue–target time intervals, i.e. whether targets appeared more likely early or late after the cue. Participants were repeatedly exposed to and trained of these environments over several days to minimize the effects of concurrent learning. Behaviorally, we found a small but robust temporal expectation effect for early intervals, which was more pronounced in naïve than in trained participants. Neurally, the expectancy effect for early intervals in trained participants was associated with significant modulations in left IPL and left RLPFC. Remarkably, when early and late likely intervals were combined, significantly enhanced fMRI-signals emerged in the left IPL, bilateral RSC, right aMPFC, and bilateral vmPFC, despite the inversion of the temporal expectation effect for late trials on the behavioral level. This neural pattern was significantly different from untrained participants especially for early time intervals in five out of six brain regions – showing expectancy effects in the trained group (for brain regions identified after extensive training). Below, we start with a discussion of the behavioral effects before turning to traditional discussion of their neural basis and end with a discussion based on brain-cognitive-term-association:

### Extensively trained temporal expectations still optimize behavior

Even with repeated exposure to the same temporal structure within a stable environment, trained participants continued to respond faster to early targets presented at likely vs. unlikely time points. The size of this temporal expectation effect was stable across runs yet smaller than a control group without prior training. The size of previously reported behavioral temporal expectation effects in single-session paradigms with comparable tasks consistently exceed the effect observed in the trained group, yet temporal expectation effects showed considerable variability across studies, ranging from over 16 ms^14,17^, to around and over 20 ms^19,44,45^, and up to more than 30 ms^46^. However, given differences in specific task designs across studies, such cross-study comparisons can only remain descriptive. Yet, the difference between trained and naive groups were also observed with the current paradigm. They may suggest that training shifted the participants’ focus from immediate history effects (see e.g.^31^) for trial history effects in foreperiod paradigms) towards the use of accumulated evidence^47^. Notably, even after extensive exposure to a stable temporal structure, temporal expectations continued to facilitate performance, indicating that well-established expectations are actively maintained and utilized to optimize behavior, consistent with the notion of everyday reliance on long-established temporal regularities.

In contrast, unlikely late trials - i.e. trials in which a stimulus does not occur after an expected delay but later – have often not been analyzed in the context of temporal research^11,48,49^; as it had been argued that these trials would contain additional cognitive processes including shifting the focus of temporal attention and reengaging to a new point in time. Moreover, some theories of attention have posited that attention only operates if there is uncertainty^50^, a condition not met by the likely late trials. Hence, effects of temporal expectation should not be observed for the late trials if expectation works by simply shifting the focus of temporal attention.

On the behavioral level, we observed responses prior to stimulus onset that varied as a function of target likelihood, consistent with the notion that temporal expectations had been formed. However, such premature responses occurred more frequently in *late unlikely* trials, resulting in shorter response times for unlikely than for likely late targets. This pattern potentially points to a failure to fully withhold responses prepared for the expected early target, thereby obscuring the corresponding response-time benefit. Moreover, it could also be argued that additional processes might have overshadowed the underlying temporal expectation effect. For instance, an expected but omitted early interval stimulus may have served as an additional cue in case of *late unlikely* trials, thereby shifting response times closer to an internally generated cue. Nevertheless, the neural signature of expectancy might even be observable for late intervals despite additional internal processes and the inverted reaction times.

### Applying overlearned temporal expectation resembles the neural pattern of retrieving episodic future simulations

To capture the neural underpinnings of overlearned temporal expectation, we initially focused on early trials only, thereby mirroring the behavioral temporal expectation effect and at the same time minimizing the contribution of potential additional cognitive processes that may have been be even more engaged during late trials. The comparison of *early likely > early unlikely* trials revealed enhanced fMRI-signals in the left IPL and left RLPFC . This finding extends previous accounts of the left IPL by suggesting that its role is not limited to the formation and implementation of newly acquired temporal expectations via temporal orienting^13,16,17,19,51^, but also encompasses the application of repeatedly used temporal expectations in stable environments. However, in contrast to the aforementioned studies, the observed effect in the present study appears to be located slightly more posteriorly, within BA 39 rather than BA 40. Nevertheless, such neuroanatomical differences should be interpreted with caution and may, at least in part, reflect variations in preprocessing and normalization procedures across studies. Unlike the left IPL, the involvement of the left RLPFC has not previously been reported in the context of temporal expectation and might therefore represent a neural signature that is more associated with overlearned rather than newly acquired temporal expectations. Modulations within RLPFC have been linked to the monitoring and evaluation of internally generated information^52,53^ and to stimulus-independent thought^54^, which becomes increasingly prominent when processing demands imposed by the external environment are low, such as in stable and predictable environments. Finally, it should be noted that previous MR-studies of temporal expectancy and attention often tested for an effect of task engagement by contrasting temporal vs. non-temporal attention (e.g.,^11^) or exogenous vs. endogenous orienting^55^, rather than investigating the effects within and across temporal environments as done here.

To assess whether temporal expectation effects generalize beyond early trials, we subsequently included late trials in the analysis. This analysis was based on the assumption that the general effect of expectancy might have been overshadowed by other processes especially for late trials on the behavioral level (see above) yet still be observable at the neural level. Indeed, we again observed a modulation in the left IPL, plus the anterior RSC, the right aMPFC, and the bilateral vmPFC for the main effect of expectancy, with the latter three regions again not previously reported in temporal expectation research.

As indexed by Neurosynth decoding results, these expectancy-modulated regions partly resemble the default mode network, which has been implicated in different higher-order cognitive processes such as self-referential cognition, mental exploration, internally generated cognition, and episodic memory processing^56,57^. The neural correlates of overlearned temporal expectation observed in the present study span two of the three proposed subsystems of the default mode network: a region of the *central core* (aMPFC) and regions of the *MTL subsystem* (IPL, RSC, vmPFC). These subsystems have distinct functions and contributions to behavioral regulation; however, during *future-oriented cognition*, these otherwise dissociable components are engaged simultaneously^58^.

Converging findings from the literature on future-oriented cognition^59–63^ have led to the proposal of a “*core network*” of the prospective brain^64^. Overlapping with the two subsystems of the default mode network previously mentioned, this network is thought to support episodic memory and episodic future simulation by temporarily decoupling from ongoing sensory input, thereby enabling the construction of internally generated representations of possible future events^65^. Accordingly, the same neural network is engaged during both episodic memory retrieval and future simulation, as both processes rely on the constructive recombination of informational content provided by episodic memory (for reviews and meta-analyses, see^66,67^. Strikingly, all expectancy-enhanced regions identified in our study fall within this “core network”, with the left IPL being causally implicated in episodic simulation by using TMS^68^, and the RSC, aMPFC and vmPFC constituting established elements of this network^67^. Accordingly, the neural correlates of overtrained temporal expectation may plausibly be interpreted within the framework of the prospective brain, which proposes that future events are mentally simulated by recombining associations acquired through past experience from episodic memory in order to generate predictions about the future^64,69^.

However, the hippocampus was not found to be engaged by overtrained temporal expectations, yet proposed to be a central component of the core network supporting episodic future thinking^59,66,70^. This could be due to the very nature of our task design. The present study involved repeatedly simulating a relatively simple and highly probable future event concerning when the target stimulus would appear rather than the construction of complex environments typically used before^71,72^. Moreover, our multi-session design may have reduced hippocampal modulations as those appear to play its strongest role during the initial construction phase when future simulations are generated for the first time, with reduced involvement once the same simulations have been repeatedly constructed^73–75^. Given repeated exposure to the same temporal environments across three consecutive days, episodic future simulations may have already been constructed during earlier sessions. Taking these findings into account, we propose that the application of overlearned temporal expectations might not only rely on the constructive recombination of episodic details that characterizes typical future simulations. Instead, overlearned temporal expectations may reflect the retrieval and reapplication of a previously established future simulation within the temporal environment in which it was originally acquired. From this perspective, overlearned temporal expectations may be conceptually related to prospective memory, that is, remembering to perform a specific action at a particular point in the future^76^, such as responding when a target is expected to occur after a particular delay.

Of course, the neural signature of overlearned temporal expectations also shares neural features with other cognitive networks, such as prospective memory, which likewise includes BA10, BA11, and IPL^77,78^, as well as memory-guided attention, which has been associated with IPL and BA10 activity^79–81^. However, the core network of episodic memory and simulation appeared to provide the best account of our findings, as it additionally includes regions implicated in future-oriented cognition, such as the RSC and vmPFC.

To avoid the fallacy of reverse inference, we directly tested whether these regions have been consistently paired not only with episodic memory but also with cognitive processes involving internally generated simulation, such as future simulation, mentalizing, and theory of mind. Using Neurosynth^37^ for both peak-wise association tests and an unbiased, data-driven decoding of the entire unthresholded second-level statistical map, we indeed observed a highly convergent functional profile. Specifically, both the peak voxel associations and the correlations with the whole-brain activation pattern were linked to terms such as autobiographical memory, episodic memory, self-referential processing, mentalizing, theory of mind, retrieval, and construction. Notably, these terms describe cognitive processes that are widely considered to rely on internally generated simulations, enabling individuals to mentally project themselves across time or into the perspectives of others, and are therefore closely related to episodic future-oriented cognition^65^. While such reverse inferences should be interpreted cautiously, the resulting cognitive profile converges on cognitive functions that have repeatedly been implicated in the prospective brain and the broader literature on episodic simulation, future-oriented and episodic cognition^38,39,65^ and is also in line with behavioral accounts linking temporal expectation with trace theories of temporal preparation^31^. These brain pattern-concept associations therefore provide convergent support for our interpretation that overlearned temporal expectations engages neural mechanisms involved in the simulation of internally represented future episodes.

### Distinct contributions of left inferior parietal lobule subregions to overtrained temporal expectations

Our findings further highlight the central role of the left IPL in overtrained temporal expectation. This region was associated with overtrained temporal expectations when additional cognitive processes were minimized by restricting the analysis to early trials and remained modulated when temporal expectations across both early and late trials were considered jointly, that is, when temporal expectations across all temporal environments were taken into account and additional cognitive processes could contribute to performance. Comparison of fMRI-modulations for trained vs untrained groups further indicate that the left IPL lobule switches preferences from *late likely* to *early unlikely* trials during training. Moreover, modulations within the left IPL were positively correlated with slower response times, suggesting that the left IPL contributes not only to the implementation of overtrained temporal expectations but also to the sustained maintenance of temporally specific action plans and the preparatory attentional states required for their execution. Although all eIects were localized within the left IPL, they formed spatially distinct clusters with only limited overlap, suggesting a measurable degree of functional heterogeneity within this region.

### No direct involvement of motor regions in overlearned temporal expectations

Temporal expectation had also been linked not only to the left inferior parietal cortex, but also repeatedly to motor regions^7,8,10,12,13,82–84^. Specifically, SMA and premotor cortex have been associated with temporal expectation not only in motor but also in perceptual tasks^85^, including rhythm prediction^86^ and in optimizing future interactions with the environment through the preparation of motor responses^85^. Importantly, motor and premotor regions have also been implicated in statistical learning of sequential regularities across a wide range of task designs^87^. This raises the possibility that the involvement of these regions in previous temporal expectation studies may partly reflect the acquisition of temporal regularities rather than the application of already established temporal expectations, as such studies have typically relied on single-session designs and averaged across trials within that session in which the observed temporal expectation effect necessarily includes ongoing learning of the underlying temporal structure. By contrast, in our multi-session design, learning might have been saturated after repeated exposure across days, as the underlying temporal structure has already become more familiar^88,89^. Consistent with this interpretation, we found no direct evidence that motor or premotor regions contribute to the application of overtrained temporal expectations. Note, however, that we used subject-specific response times as a covariate and observed variations associated with this explanatory variable in the left primary motor cortex, SMA, and pre-SMA, suggesting a role in faster responding.

### Temporal unexpectedness

As expected, temporally unlikely targets elicited activity in a predominantly right-lateralized network including the temporoparietal junction, anterior insula, visual cortex, premotor cortex, and inferior frontal gyrus, among other regions. This pattern closely resembles networks previously associated with attentional reorienting and the detection of behaviorally relevant cross-modal changes in the environment^42,43^, thereby supporting the validity of our temporal expectation manipulation. Within a predictive coding framework, these activations may reflect processes involved in the detection of violations of (temporal) predictions and the subsequent updating of internal models. In particular, the recruitment of the right anterior insula together with the right inferior frontal gyrus (pars opercularis, BA44), is consistent with a salience network that signals the occurrence of task relevant or unexpected stimuli and initiates cognitive control processes. The concurrent involvement of primary motor cortex, the SMA and pre-SMA, as well as premotor cortex, further suggests the engagement of mechanisms supporting the rapid updating and adjustment of action plans in response to unexpected events. In addition, right-lateralized supramarginal gyrus activity might reflect attentional reorienting to unexpected stimuli . Enhanced fMRI-signals in visual cortices likely reflects enhanced sensory responses to unexpected stimuli, a well-established effect in accord with predictive coding. All these regions appeared to remain responsive to temporal unlikeliness after excessive training, suggesting that these regions contribute to the detection of expectancy violations even in an overtrained state rather than to the implementation of established temporal predictions. The enhanced involvement of inferior frontal and temporoparietal regions especially for late trials may indicate that additional processes including temporal reallocation, surprise ad updating might be computed there. This interpretation is in agreement with recent fMRI-studies focusing on expectation violation which related the right temporoparietal junction to attentional reallocation using fMRI and EEG^43^; while others highlighted the role of frontal and parietal regions in surprise and updating^51^.

Within the framework of predictive coding it has been proposed that putatively enhanced neural signals for unexpected relative to expected trials may reflect a dampening of expected rather than the enhancement of unexpected signals; though response sharpening as a function of expectation has also been observed^90^. In our study a more nuanced picture emerged regarding the neural underpinnings of temporal expectations outside sensory areas. First, we identified a consistent network of regions instrumental in the application of overlearned expectations: in left parietal areas overlearned expectations lead to increased expectancy-related fMRI-signals for early intervals together with stable expectancy-driven response for late intervals potentially suggesting response sharpening in this region. In frontal regions overall response dampening could be observed over time together with an increase in the differential expectancy-related signal. This pattern of results evolves over time and thereby appears to maintain and utilize temporal expectations. Future studies are needed to differentiate to which extent this emerging brain network is driven by passive observation, active training or a mixture of both.

### Conclusions

Taken together, our findings indicate that the neural underpinnings of temporal expectations differ as a function of training status as the involvement of training-related brain regions may only become apparent once temporal expectations have been established and repeatedly utilized over extended periods of time. We further observed that brain regions commonly associated with episodic context-dependent future simulations are used to maintain and apply even simple expectations. This raises the possibility that learned cue–target intervals constitute a minimal representational scaffold of an episode. Rather than requiring rich scenes, narratives, or autobiographical content, the neural machinery commonly associated with episodic cognition may already be engaged by the encoding and retrieval of simple temporal relations between events. These cue-triggered recalled simulations may serve as a basis for predictions that guide goal-directed actions and optimize behavior.

## Methods

### Participants

Participants were recruited at the Otto-von-Guericke University Magdeburg, Germany. The study protocol was approved by the local ethics committee. All participants provided informed written consent before participation and received monetary compensation for taking part in the study. Only participants were included if they reported no history of psychiatric or neurological disorder, no regular intake of medication known to interact with the central nervous system, normal or corrected-to-normal vision and no hearing impairments.

Forty-two healthy volunteers participated in the study. From the final analysis, six participants were excluded due to technical issues during MRI data collection and one additional participant was excluded due to extreme parameter estimates identified during quality-control procedures, resulting in 35 participants (19 female, mean age = 26.4 y +/-5.2 standard deviation (SD)). For the control study, data from both an MRI cohort and an additional behavioral-only cohort were combined. Both cohorts completed an identical task using the same experimental procedure, differing only in the testing setting (inside vs. outside the MRI scanner). The MRI cohort comprised 16 participants (10 female, mean age = 23.3 y ± 4.5 SD), whereas the behavioral-only cohort comprised 35 participants (26 female, mean age = 22.4 y ± 3.3 SD), resulting in a total control sample of 51 participants (36 female, mean age = 22.3 y +/- 3.7 SD),

### Experimental procedure

Our participants repeatedly performed two versions of a task across 5 sessions on 4 consecutive days in the main experiment (Fig. 1). The control experiment was identical to the main experiment except that participants received no prior training or observations (no session 1-4) and only performed one active session, either inside the MR-scanner (identical to session 5 in the main experiment) or outside the MR-scanner (identical two sessions 2-3 outside the scanner). On day 1 participants passively observed auditory and visual cue-target combinations inside the MRI-scanner (session 1; *passive naive*), unaware of what would happen on the following days or the relational structure between the stimuli. On day 2 and 3 they were trained on visual target detection outside the MRI-scanner (session 2 and 3; *active training*) and on day 4 they first passively observed the cue-target parings (session 4; *passive trained*) before they actively detected visual targets again inside the MRI-scanner (session 5; *active trained*). In the present article, we focus exclusively on session 5; analyses of the other MRI sessions are beyond the scope of the article and will be reported elsewhere. In all sessions, visual stimuli appeared at one of two possible time points after an auditory cue: either early (500 ms) or late (1500 ms). The likelihood of these two points in time was manipulated in a block-wise manner, with one block favoring early onsets (early:late ratio of 4:1) and the other favoring late onsets (early:late ratio of 1:4), see Fig. 1. These two block types (50 trials per block) alternated throughout the experiment. Within each block, one time interval was more likely (e.g., early in the early-likely block), while the other time interval was unlikely (e.g., late in the early-likely block). This block-wise manipulation established two distinct temporal environments: one in which early target onsets were more probable and another in which late target onsets were more probable. By structuring the experiment in this manner, we created an adaptive environment in which participants could form context-dependent temporal expectations based on the probabilistic structure of each environment. Participants were not informed at any point about the temporal relationship between the auditory and visual stimuli, nor were they explicitly told that the auditory stimulus could be used as a cue to predict the visual target.

### Stimuli and task

Each session consisted of four runs, with 100 trials per run. Within every run, both the early-likely and late-likely blocks were presented, each containing 50 trials. In these blocks, the target appeared at the likely time point in 37 trials and at the unlikely time point in 13 trials. A trial began with the auditory cue (presented for 100 ms), followed by the target (also presented for 100 ms) at either the early or late time point (see Fig. 8). The starting block was counterbalanced across participants.

**Fig 8.**
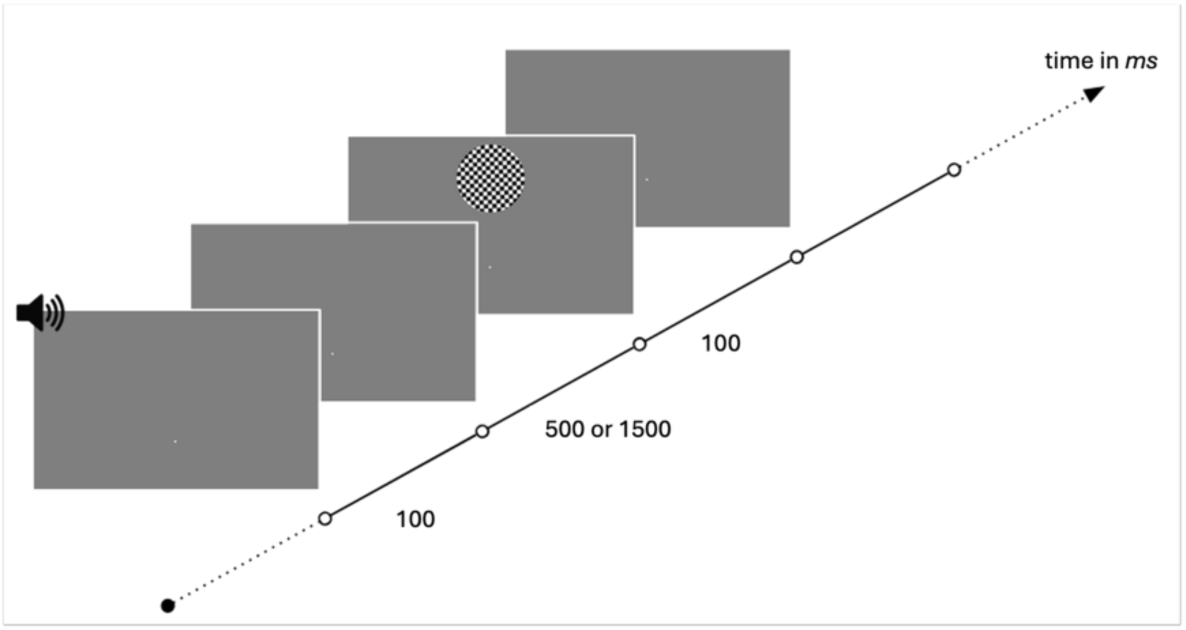
Trial structure. Each trial began with an auditory cue (100 ms), followed after either 500 ms or 1500 ms by a visual target (100 ms).

Participants were asked to maintain fixation at all times, both during the passive and active sessions. In the active task, they were also instructed to respond as quickly as possible by pressing a button when they detected the target stimulus (see Fig. 8).

In case a participant did not respond within 800 after target appearance, the response was considered missing. The following trial began after an inter-trial interval drawn from a truncated exponential distribution (minimum 0.5 s, maximum 8 s, discretized in steps of 0.1 s). For the main MRI-experiment, a predefined set of inter-trial intervals was used (based on design efficiency optimization^91^). Both the cue-target time interval (early or late) of a trial and the block type (*early likely* or *late likely*) together determined whether the target was presented as early-likely, late-unlikely, late-likely, or early-unlikely.

All stimuli were created and presented using the Psychophysics Toolbox ^92^ within MATLAB 2012b (MathWorks, Inc.) running on a Windows 7 environment. Visual targets consisted of circles filled with a checkerboard pattern (∼10.3° of visual angle, 400 pixels diameter, changing from black to white every 25 pixels) and were centered in the upper third of the screen. Visual stimuli were displayed either via an LCD projector (resolution 1920 x 1080 pixels, frame rate = 60 Hz) on a screen placed at the rear opening of the bore of the MRI scanner, and viewed through a mirror mounted on the head coil (distance to screen = 35 cm) or via an LCD screen (22” 120 Hz, SAMSUNG 2233RZ) with adjusted resolution to 1920 x 1080 pixels and the refresh rate of 60 Hz in the behavioral laboratory (distance to screen = 103 cm). Auditory stimuli were clearly audible pure tones (8000 Hz) and presented via headphones (Sennheiser HD 650 in the behavioral laboratory, mark II-Box; MR CONFON in the scanner). The experiment had a dark grey background (RGB: 125) with a white fixation square (20 pixels) centered in the lower third of the screen. Participants responded with a wireless mouse (Logitech M325; behavioral laboratory) or with an MRI-compatible response box (Lumitouch; MR scanner).

### Behavioral data analysis

The behavioral analyses examined how response times varied for likely and unlikely presented targets, with a specific focus on early targets^48,93^ to assess the temporal expectation effect on response times (e.g. shorter response times for likely presented targets compared to unlikely ones); and further whether effects would change with training status in accord with ^8^.

For the statistical analysis, we first used preplanned t-tests to investigate the difference between *early likely* and *early unlikely* trials. In addition, we also performed a repeated measures ANOVA with likeliness (likely, unlikely) and run (1, 2, 3, 4) as within-subject factors to analyze the robustness of this effect over time. The same repeated measures ANOVAs were subsequently performed for late targets and the effects in the untrained group. Greenhouse-Geisser corrections were applied where necessary.

To investigate the effect of training status, we used a linear mixed-effects model due to the unequal sample sizes between training groups and the presence of incomplete cells in the design. For early targets, response times were modeled as a function of likeliness, training status, and their interaction, with laboratory (inside, outside the scanner) included as an additional fixed effect. Subject was included as a random intercept:

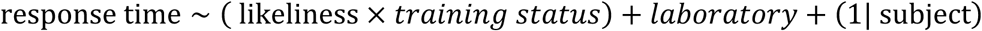

In order to correct for multiple comparisons, FDR correction was applied. A significance level of p < 0.05 was used to determine statistical significance. The behavioral analyses were performed using R Statistical Software (v4.1.2; R Core Team 2021) using the package afex^94^.

### Neuroimaging data acquisition and preprocessing

Magnetic resonance imaging was conducted on a Siemens Prisma 3 Tesla system (Erlangen, Germany) with a 64-channel head coil for signal reception. T1-weighted structural images were acquired with an MPRAGE sequence using the following parameters: 1 × 1 × 1 mm3 voxel size, 256 × 256 × 192 matrix, 2.82 ms echo time, 2.5 s repetition time, 1.1 s inversion time, 7° flip angle, 140 Hz per pixel bandwidth, 7/8 partial Fourier, parallel imaging with a GRAPPA factor of 2, and 5:18-min scan duration. For functional imaging we used a multiband accelerated T2*-weighted echo-planar imaging sequence (multiband acceleration factor 2, repetition time = 2000 ms, echo time = 30 ms, flip angle = 80°, field of view = 220 mm, voxel size=2.2×2.2×2.2 mm, and no gap). During each of the functional runs we acquired 250 volumes. Per volume, 66 slices covering the whole brain, tilted by approximately 15° in z-direction relative to the anterior–posterior commissure plane were acquired in interleaved order. Data preprocessing was conducted using fMRIPrep 21.0.2^95^. T1-weighted structural images were corrected for intensity non-uniformity, and used as T1w-reference throughout the workflow. The T1w-reference was then skull-stripped, segmented and normalized to the Montreal Neurological Institute (MNI 152) template. For functional imaging preprocessing, BOLD runs were slice-time corrected to the middle slice, were corrected for head-motion by applying beforehand estimated head-motion parameters and were then co-registered to the T1w reference. Resulting images were spatially normalized to MNI-space using ANTs’ *antsRegistration* in a multiscale, mutual-information based, nonlinear registration scheme and were then smoothed with a Gaussian kernel of 8 mm full-width at half maximum (FWHM) via SPM12 (Wellcome Trust Centre for Neuroimaging, London, UK).

### Neuroimaging data analysis

For the statistical analysis of the fMRI data, we time-locked the onsets on the cue presentation to capture brain signals related to target expectation. The goals of the neuroimaging analyses were to identify brain areas linked to the differentiation of likely and unlikely events in the distinct temporal environments (time interval x likeliness). The first-level GLM was constructed, utilizing the hemodynamic response function (HRF), time derivatives (TD), and dispersion derivatives (DD) to model the BOLD signal. The model assessed the neural response to all four experimental conditions: *early likely*, *early unlikely*, *late likely* and *late unlikely*. This first-level GLM included those experimental conditions as predictors for all collected sessions combined. In addition, we included six movement parameters (estimated during realignment) per run as nuisance regressors. To assess the mean activity per condition, contrasts averaging across the four runs for each session were constructed on the first-level, and a session-wise flexible factorial with those mean activity per condition and additionally mean response time per condition and subject was performed on the group level. All analyses were performed using the canonical hemodynamic response function (HRF) using SPM12 (Wellcome Trust Centre for Neuroimaging, London, UK). All whole-brain results were thresholded at p < .001 (uncorrected) at the voxel level and corrected for multiple comparisons at the cluster level using family-wise error (FWE) correction (p < .05). Marsbar ^96^ was used for mean beta weight extraction. Post hoc tests were conducted using R Statistical Software (v4.1.2; R Core Team 2021) using the package afex^94^. fMRI-preprocessing and analysis of the fMRI-control study was identical to the main experiment. Due to the lower number of participants in the control study, the result maps from the critical comparison of the main experiment (likely > unlikely) were used as masks. Region-wise mean beta weights were extracted and analyzed using R (v4.1.2; R Core Team 2021).

To assess whether the identified peak voxels for the contrast *likely > unlikely* corresponded to regions consistently associated with episodic cognition, we queried the Neurosynth^37^ association map for the term *episodic*. For each peak voxel, the corresponding association z-score was extracted from the meta-analytic map. To further characterize the overall cognitive profile of the observed activation pattern, the unthresholded group-level statistical map for the contrast *likely > unlikely* was decoded using the Neurosynth Decoder^37^ via NeuroVault^97^. The decoder quantifies the spatial similarity between the statistical map and meta-analytic activation maps associated with cognitive terms.

## Data availability statement

All behavioral data and the fMRI contrast images used for the present analyses are publicly available via OSF at https://osf.io/am6y7/. Owing to the OSF storage limit (50 GB), the repository includes the complete preprocessed fMRI dataset for the naïve control group and a subset of the preprocessed fMRI data for the trained group. Researchers interested in accessing the de-identified raw MRI data or the complete preprocessed fMRI dataset (approximately 80 GB of additional data) for non-commercial research purposes are invited to contact Linda Sempf with a brief description of their intended use. Access will be provided upon review of the request. Code availability: All code used for the behavioral data analyses, fMRI preprocessing, first-level analyses, and second-level group analyses is publicly available on GitHub at https://github.com/lsempf/OTE-sfb1436.

## CRediT authorship contribution statement

**Linda Sempf**: Writing – original draft, Visualization, Methodology, Investigation, Formal analysis, Data curation, Conceptualization. **Peter Vavra**: Writing – original draft, Methodology. **Toemme Noesselt**: Writing – original draft, Methodology, Supervision, Funding acquisition, Conceptualization.

